# 2-photon-fabricated nano-fluidic traps for extended detection of single macromolecules and colloids in solution

**DOI:** 10.1101/2021.11.17.468989

**Authors:** Oliver Vanderpoorten, Ali Nawaz Babar, Georg Krainer, Raphaёl P.B. Jacquat, Pavan K. Challa, Quentin Peter, Zenon Toprakcioglu, Catherine K. Xu, Ulrich F. Keyser, Jeremy Baumberg, Clemens F. Kaminski, Tuomas P. J. Knowles

## Abstract

The analysis of nanoscopic species, such as proteins and colloidal assemblies, at the single-molecule level has become vital in many areas of fundamental and applied research. Approaches to increase the detection timescales for single molecules in solution without immobilising them onto a substrate surface and applying external fields are much sought after. Here we present an easy-to-implement and versatile nanofluidics-based approach that enables increased observational-timescale analysis of single biomacromolecules and nanoscale colloids in solution. We use two-photon-based hybrid lithography in conjunction with soft lithography to fabricate nanofluidic devices with nano-trapping geometries down to 100 nm in height. We provide a rigorous description and characterisation of the fabrication route that enables the writing of nanoscopic 3D structures directly in photoresist and allows for the integration of nano-trapping and nano-channel geometries within micro-channel devices. Using confocal fluorescence burst detection, we validated the functionality of particle confinement in our nano-trap geometries through measurement of particle residence times. All species under study, including nanoscale colloids, α-synuclein oligomers, and double-stranded DNA, showed a three to five-fold increase in average residence time in the detection volume of nano-traps, due to the additional local steric confinement, in comparison to free space diffusion in a nearby micro-channel. Our approach thus opens-up the possibility for single-molecule studies at prolonged observational timescales to analyse and detect nanoparticles and protein assemblies in solution without the need for surface immobilisation.

## Introduction

The spatial confinement of biomolecules or colloidal nanoparticles in solution for biophysical studies at the single-molecule level has become instrumental in many areas of fundamental and applied research including nanobiotechnology [1], biophysics [2], and clinical diagnostics [3]. It allows for increased observational-timescale analysis of nanoscopic species such as nucleic acids, protein assemblies [4] or colloidal particles [5] with single-molecule sensitivity [6]. Currently, molecular confinement is most typically achieved through surface immobilisation of the biomolecule or nanoparticle of interest on a substrate surface (e.g., for confocal or total internal reflection fluorescence (TIRF) microscopy) [7],[8],[9]. This approach, however, has numerous drawbacks, not least because surface interactions can change the molecule’s configuration and function.

An alternative to surface immobilisation is the trapping of particles in solution without immobilising them onto a substrate surface. Various approaches using external fields, such as electric [10], hydrodynamic [11] and optical fields [12], [13], for nanoparticle trapping in solution have emerged. Optical trapping, for example, has proven effective in measuring repulsive or attractive forces between particles such as colloids and proteins, but the high laser powers required induce flows around the trapped particles leading to undesirable and confounding effects [1]. Furthermore, such techniques suffer from low throughput and require a refractive index mismatch between the particle and its surrounding media [14], which is often not the case when monitoring biological specimens. Other techniques, such as thermal trapping [15]–[17], have also shown to be effective at confining nanoparticles in small volumes, but similar to optical trapping, thermal particle trapping has significant drawbacks due to the sample undergoing motion because of convection. This puts limitations on the estimation of particle properties such as molecular size and particle reaction kinetics at physiologically relevant conditions.

Recently, geometry-induced electrostatic trapping and colloidal trapping based on the spatial modulation of configurational entropy was demonstrated [18],[19]. This approach enables trapping without applying external fields and has proven invaluable in observing particles in an all aqueous environment [20]. Mojarad et al. [21] demonstrated trapping of colloids and gold nanoparticles in nanofluidic silica devices, which allowed measurement of their particle size and charge in silica-based nano-wells. Ruggeri et al. [22],[23] further pushed the limits of nano-trapping-based electrometry to the single-molecule level. While efficient in their use, however, to date, the fabrication of such trapping devices and their subsequent integration with microfluidic device platforms is challenging and demands specialised clean room equipment such as electron beam lithography (EBL) [24] and reactive ion etching (RIE) [25]. Even though such approaches generate nano-slits or nano-channels smaller than 100 nm [26], the complexity of the fabrication process, writing times, and the costs to produce a single device render these techniques highly inefficient and impractical. Additionally, most of these techniques are relatively low throughput and integrating them with micro-channels, which is required for the chip-to-world interface, can be challenging.

An alternative approach for fabricating nano-traps/nano-channels and integrating the nanostructures within a microfluidic chip platform involves the combination of conventional UV lithography followed by two-photon lithography (2PL) [27],[28], where a focused femto-second pulsed laser is scanned across the photoresist, resulting in the writing of device features below 200 nm in lateral size. 2PL or direct laser writing (DLW) is a powerful emerging technology and has gained much attention in the last years for the fabrication of 3-dimensional (3D) micro- and nano-structures and functional devices below the diffraction limit [29]. Fabrication of arbitrary 3D structures is possible in a photoresist from computer-generated 3D models and thus constitutes a fast and straightforward fabrication procedure [30]. Previously, microfluidic [31], nanofluidic [32], and optofluidic [33] devices were fabricated using femto-second laser 3D micromachining and were shown to allow for the integration of functionalities unachievable with conventional UV-lithography in device designs.

Here, we demonstrate the facile fabrication of nanofluidic trapping devices using a 2PL system for increased observational-timescale single-molecule studies of biomacromolecules and colloids in solution. To this end, we developed an approach based on hybrid 2PL- and UV-lithography in conjunction with soft lithography [34] to generate nanoscale channels and adjacent nanoscale trap structures with dimensions down to 100 nm in height in a single step from a silicon master wafer. This allowed for the fabrication and prototyping of nanofluidic polydimethylsiloxane (PDMS)–silica devices in a facile and scalable manner and the writing of various nano-trapping geometry designs with varying heights in one writing process. We analysed the master wafer and PDMS imprints using correlative scanning electron microscope (SEM) and atomic force microscope (AFM) characterisation techniques and validated the functionality of particle confinement in nano-trap geometries through measurement of particle residence times in nano-traps as compared to micro-channels and nano-channels using single-molecule fluorescence burst analysis. We found that all species analysed, including nanoscale colloids, protein oligomers, and short DNA duplexes, showed a three- to five-fold increase in average residence time in the detection volume of nano-traps in comparison to free space diffusion in a nearby nano- or micro-channel. We further demonstrate other fluorescence microscopy techniques (confocal imaging and TIRF microscopy) as alternative readout techniques to be used in combination with nanofluidic traps. Taken together, our developments presented herein constitute a cost-effective and easy-to-implement approach for the fabrication of nanofluidic trap devices and open-up a broad avenue of possibilities to study single molecules in solution for extended periods of time without permanent surface immobilization and without applying external fields.

## Results and Discussion

### Integration of nano-trapping and nano-channel geometries between micro-channels with 2-photon lithography

Conventional fabrication of trapping devices relies on sophisticated clean-room equipment [18] and does not allow high throughput and flexibility in the writing of structures of varying geometry and height. To overcome these challenges and make the fabrication process more facile, we propose here a fabrication route of nanofluidic devices via hybrid 2PL that enables the writing of nanoscopic 3D structures directly in photoresist [28]. By combining large area UV mask lithography with local high precision two-photon laser writing, we demonstrate the integration of nano-traps written adjacent to nano-channels in a pre-existing microfluidic device design (see **Figure 1**). Since 2PL is a dosage-dependent process and the smallest feature size obtained in the photoresist depends on the laser intensity and exposure time, we first set out to first optimise the fabrication procedure to achieve full merging of nano-trap and nano-channel geometries.

**Figure 1.**
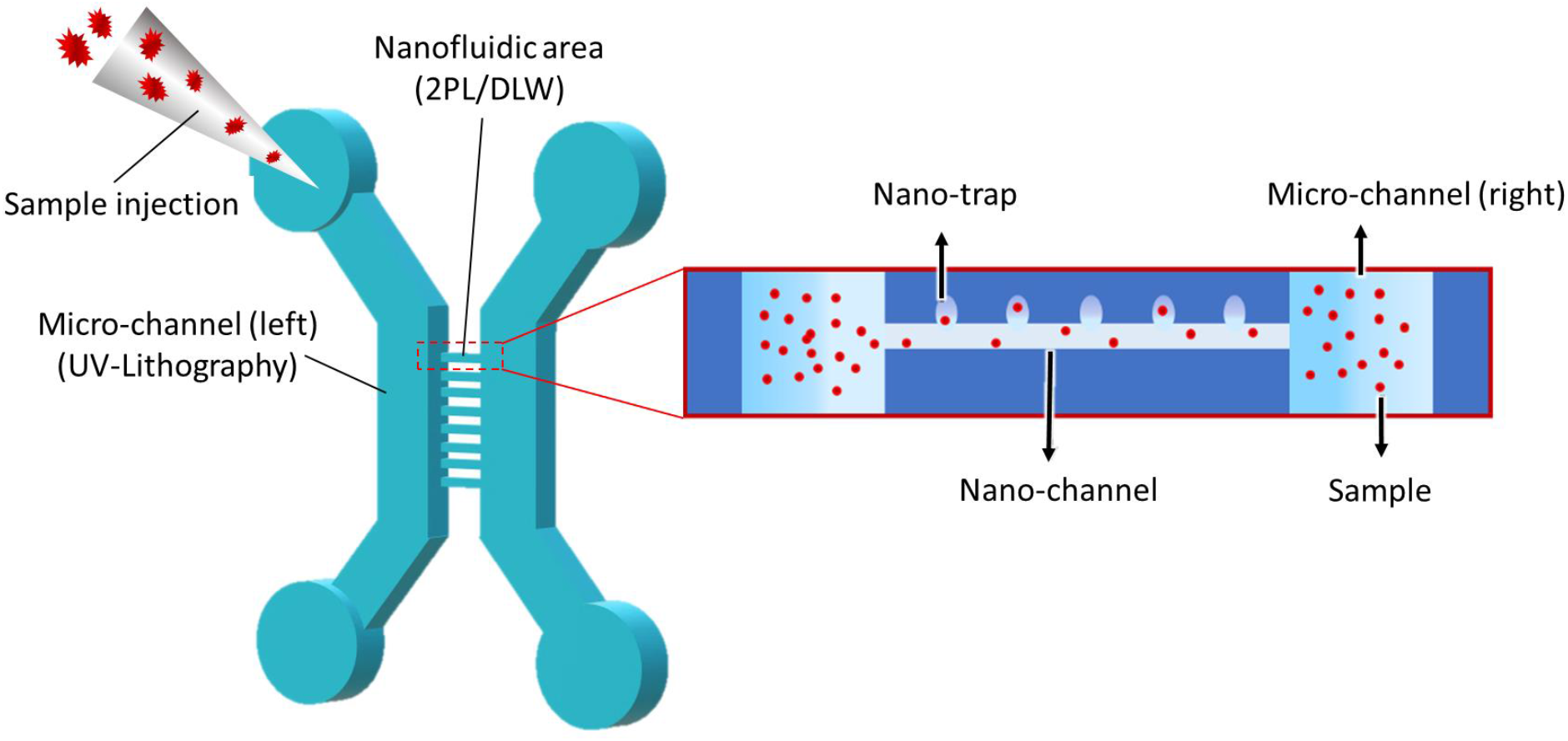
Design and fabrication of nanofluidic device with trapping functionalities. Schematic of the device design consisting of microfluidic reservoirs, inlets/outlets, nanofluidic channels and nano-trapping arrays. 2-photon lithography (2PL) (or direct laser writing, DLW) is used to combine microfluidics with nanofluidic functionalities. Large area mask-based UV lithography patterns microfluidic areas, whereas 2PL incorporates nano-channels and nano-traps in between two micro-channels. The inset illustrates the placement of the nano-traps next to the nanofluidic channel.

We began by exploring and prototyping nanofluidic geometries in negative SU-8 photoresist (**Figure 2**) and produced imprints into PDMS following standard UV- and soft-lithography protocols (**Figure 3**). Characterization techniques such as SEM and AFM were used to analyse the prototype nanostructures. Varying the laser power, laser writing speed, and the distance in between the nano-traps and nano-channel (Y-offset) during the 2PL writing process resulted in different configurations of nano-trap moulds as shown in **Figure 2 (A-E)**. Straight nano-channels were written at a fixed laser intensity of 90 mW and a writing speed of 100 μm/s. Dots for nano-trap moulds were written adjacently with 1000 μm/s scanning speed and by modulating the laser at 100 mW. Nano-traps were added every 3 μm along the nano-channels. The height of the nano-traps was smaller than the nano-channels due to the lower net exposure of the photoresist. Notably, the 3D piezo-flexure stage used for scanning of the laser beam is a key component and allowed for varying the Y-offset between nano-traps and nano-channels with a resolution down to 10 nm by leveraging the closed-loop control mode of a piezo stage. Accordingly, the Y-offset was varied from 1.25 μm to 0.85 μm in steps of 100 nm. As shown in **Figure 2 (B)**, at a Y-offset of 1.15 μm, the SU-8 of the nano-trap geometry merged with the nano-channel through monomer cross-linking. The same geometries were also analysed in the PDMS imprints as shown in SEM micrographs of **Figure 3 (A-E)**. Notably, by just varying the Y-offset between the nano-channel and nano-traps, different geometries and designs of the nano-traps in PDMS could be generated, for example, triangular nano-traps as shown in **Figure 3 (B)**. This highlights the importance of precise laser positioning to control not only the merging of nano-channels with nano-traps, but also the possibility to create traps with varying geometries. The process of 2PL for writing almost arbitrary 3D structures thus allows significant flexibility here for choosing and modulating the desired geometry, microfluidic chip design, and introducing multiple geometry layers within a single spin-coating process. Indeed, we were able to add other conformations of traps to a nano-channel, for example, where the traps were positioned on top of the nano-channels (**Figure 3 (G)**) or nano-traps with bottle-neck openings (**Figure 3 (H)**) on the side. The latter structures exhibited a nano-trap height of 100 nm, as confirmed by correlative SEM/AFM measurements on the master wafer (**Supplementary Figure 1**).

**Figure 2.**
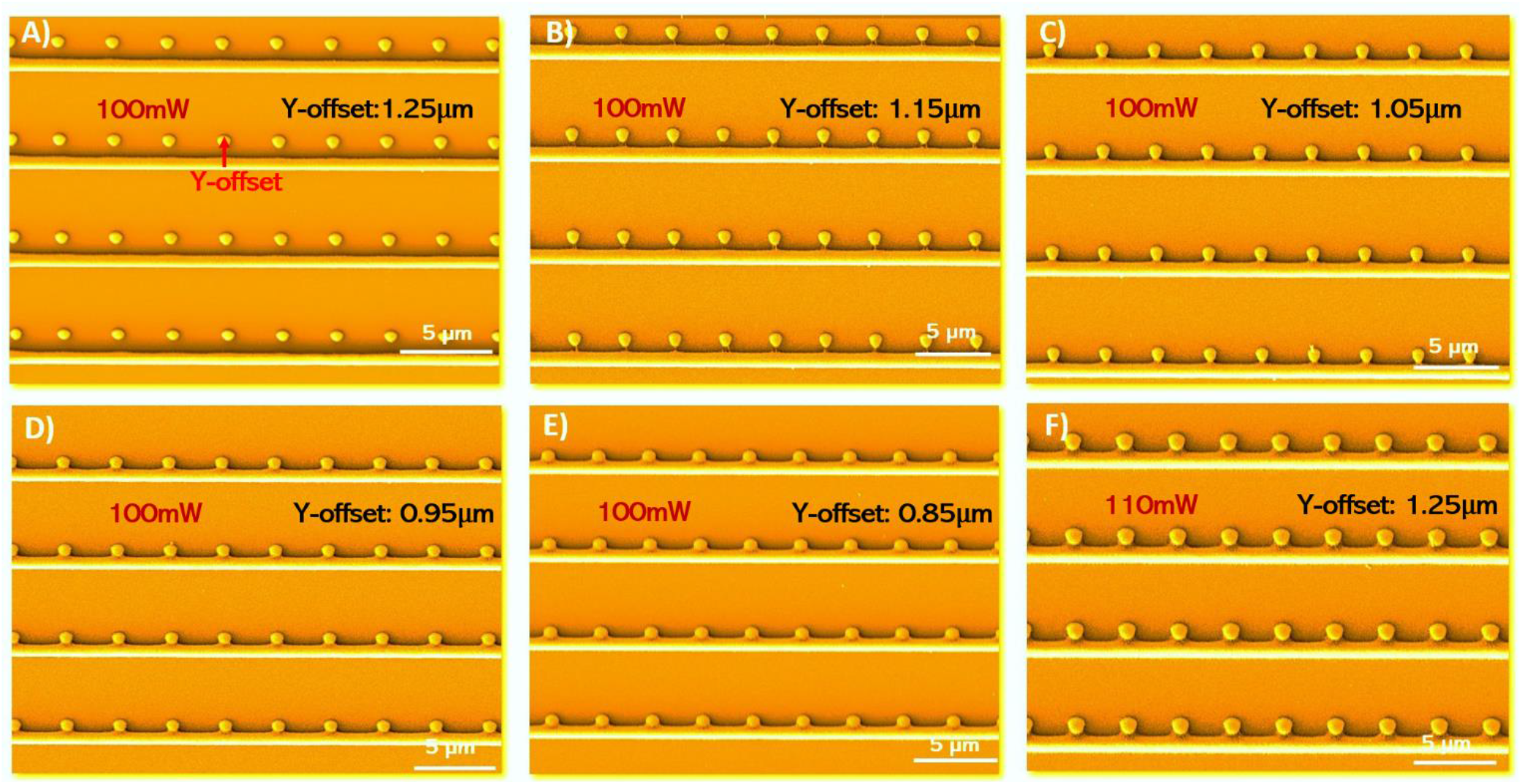
Prototypes of nano-channel and nano-trap geometries fabricated in photoresist using 2-photon lithography. Shown are SEM micrographs of nano-channel/nano-trap moulds as obtained by 2-photon lithography in SU-8 photoresist using varying laser powers and Y-offsets. The writing speeds for the nano-traps and nano-channels were 1000 μm/s and 100 μm/s, respectively. **(A)–(E)** Nano-channel/nano-trap moulds obtained with a Y-offset in the range of 1.25–0.85 μm; the laser power for writing nano-channels and nano-traps was 90 mW and 100 mW, respectively. **(F)** Optimized nano-channel/nano-trap mould written with a Y-offset of 1.25 μm; the laser power for writing nano-channels and nano-traps was 100 mW and 110 mW, respectively.

**Figure 3.**
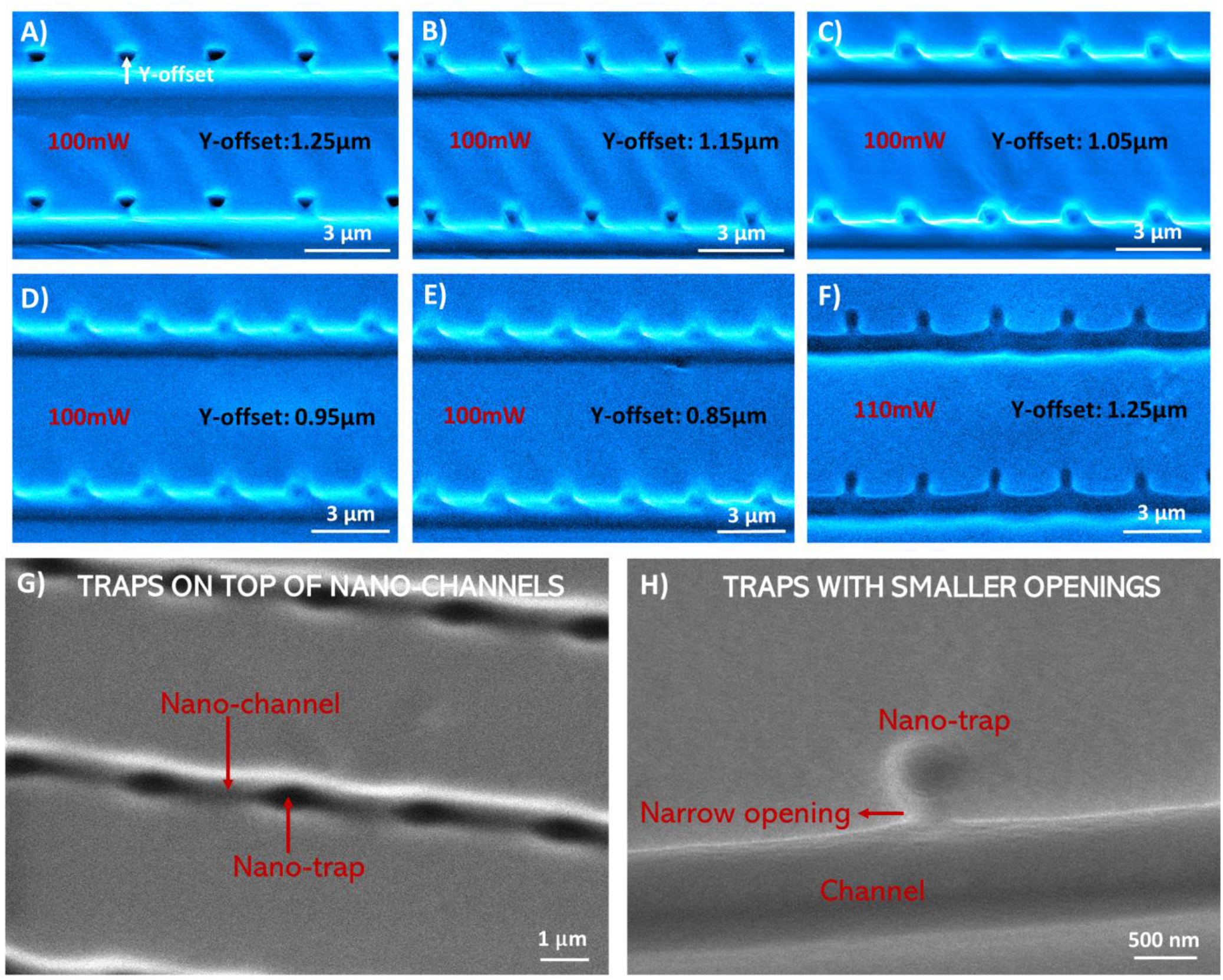
PDMS imprints of nano-channel and nano-trap device prototypes. Shown are SEM micrographs for nano-channels and nano-traps imprinted in PDMS. The moulds, from which the PDMS imprints were fabricated, were written in SU-8 photoresist with 2PL by varying the laser power and Y-offset (Figure 2). The writing speeds for the nano-traps and nano-channels were 1000 μm/s and 100 μm/s, respectively. **(A)–(E)** Nano-channels and nano-traps imprinted in PDMS with Y-offset in the range from of 1.25 μm–0.85 μm; the laser power for nano-channels and nano-traps were 90 mW and 100 mW, respectively. **(F)** Optimized nano-channel/nano-traps imprinted in PDMS with Y-offset of 1.25 μm; the laser power for nano-channels and nano-traps were 100 mW and 110 mW, respectively. **(G)** SEM image of a trapping device with nano-traps on top of nano-channels in the PDMS (top view). **(H)** SEM image of the narrow opening of a nano-trap imprinted in PDMS. Correlative AFM imaging showed a height of approx. 100 nm of the pockets (**Supplementary Figure 1**).

The prototyping geometries obtained thus far were used to determine appropriate and optimised writing parameters for creating nanofluidic trapping devices required for nanoparticle and biomolecule trapping in single-molecule experiments (see below). For this chip design, we required round nano-trapping cavities of a few hundred nanometres in radius which are well-merged with straight nano-channels that have dimensions in the submicron-regime. Such geometrical features could be obtained by using 110 mW laser power for writing of the traps, 100 mW for the nano-channels and a Y-offset in between them of 1.25 μm (**Figure 2 (F)**). Thereby, we fabricated nano-traps of 350 nm in radius adjacent to nano-channels of 650 nm in width. The chosen fabrication parameters show geometrical consistency between individual traps and are still mechanically stable enough to have the same structures in the final bonded device. The mechanical stability of the nano-trap structures in SU-8 was further enhanced by increasing the cross-linking density of monomers with a second UV exposure after writing nanostructures with 2PL [35].

### Integration of nano-channel and nano-trap geometries in a microfluidic device platform

After having optimised the procedures for generating nano-trap and nano-channel geometries via our 2PL approach, we set out to fabricate the combined nanofluidic device for single-molecule experiments, as shown in **Figure 1**. The device was produced by first generating the micron-scale structures of the chip, which consisted of two microfluidic channels and reservoirs, sample inlets/outlets and pre-filters. This was done by transferring these chip features from a high-resolution transparency acetate photomask onto SU-8 photoresist, spin-coated on a silicon wafer, via conventional contact UV lithography [30]. In a second step, the microfluidic channel reservoirs, separated by 75 μm, were connected with straight nano-channels and adjacent nano-traps using the optimised 2PL writing parameters, as detailed above (c.f., **Figure 2 (F)** and **Figure 3 (F)**). Subsequently, PDMS imprints and glass-bonded chips were produced from these structures using standard soft lithography and replica moulding procedures. **Figure 4 (A)** shows a SEM micrograph of the final PDMS imprint with an overview of the conventional micron-scale chip functionalities. Further magnification (**Figure 4 (B)–(D)**) shows the successful integration of nanofluidic functionalities in between the microfluidic reservoirs. Two microfluidic compartments of 25 μm depth were joined by 2PL with six nanofluidic areas (**Figure 4 (B)**, indicated with arrows). **Figure 4 (C)** shows in greater detail one nano-trapping array consisting of 18 nano-channels with adjacently added nano-traps every 3 μm. Notably, the channels show a wider funnel-like shape at the microfluidic interface due to the sequential double exposure of the photoresist by UV-lithography and 2PL. The central part of the array, however, shows the intended trap geometry from the prototypic procedure above, with suitable traps for confinement of nanoparticles imprinted in PDMS. The nano-channels were 650 nm wide and connected to the nano-traps, which had a radius of 350 nm. The nano-channels and nano-traps were 750 nm and 650 nm in height, respectively, according to correlative profilometer measurements (**Supplementary Figure 2**).

**Figure 4.**
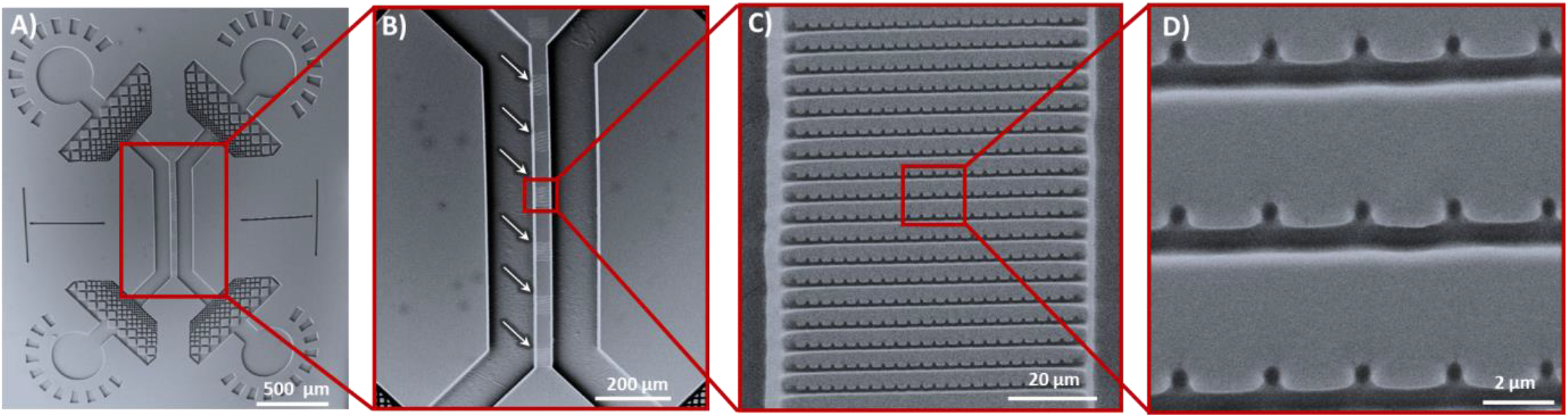
Nanofluidic device with trapping functionalities for single-molecule experiments. Shown are SEM micrographs of PDMS nanofluidic device imprints fabricated via hybrid UV mask lithography and 2PL. **(A)** Full view of the micro- /nanofluidic device, consisting of microfluidic reservoirs, inlets/outlets, nanofluidic channels and nano-trapping arrays. The design corresponds to the schematic shown in Figure 1. **(B)** Magnification depicting the arrays of 75 μm long nano-channels with integrated nano-traps in between the two 25 μm deep micro-channels in PDMS. **(C)** Higher magnification of nanofluidic channels and nano-traps shows consistent imprinting of nano-trapping arrays in PDMS. **(D)** Zoom-in of SEM micrograph showing the geometry of nano-traps.

### Single-molecule fluorescence detection of colloids and biomolecules in nano-traps

Single-molecule studies for biological measurements in miniaturised devices have proven very useful due to their precise sample handling, small volume manipulation, and high throughput capabilities [36], [37]. Prolonged observation of single molecules or nano-colloids in solution is still a challenging task but an important step towards microfluidic total-analysis systems (μTAS).[38] Our chip design provides an opportunity for prolonged detection of single particles in solution without permanent surface immobilization. We intend to increase particle residence times in a detection volume due to the additional local steric confinement in the nano-traps.

To demonstrate this, we set out confocal-based single-molecule burst experiments that allowed us to observe, record, and compare the events of single particles entering and leaving the nano-trapping geometry. **Figure 5** schematically illustrates the experimental setup. The device’s micro-channel reservoirs were filled with respective particle solutions at pico-to nano-molar concentrations. Once the sample in the device reached equilibrium and the nanoparticles started diffusing through the nano-channels, fluorescence burst detection was conducted within the nano-traps. Samples were excited with a continuous 488-nm diode laser and their fluorescence collected using avalanche photodiodes, which allowed readout of the fluorescent nanoparticle signal with high temporal resolution.

**Figure 5.**
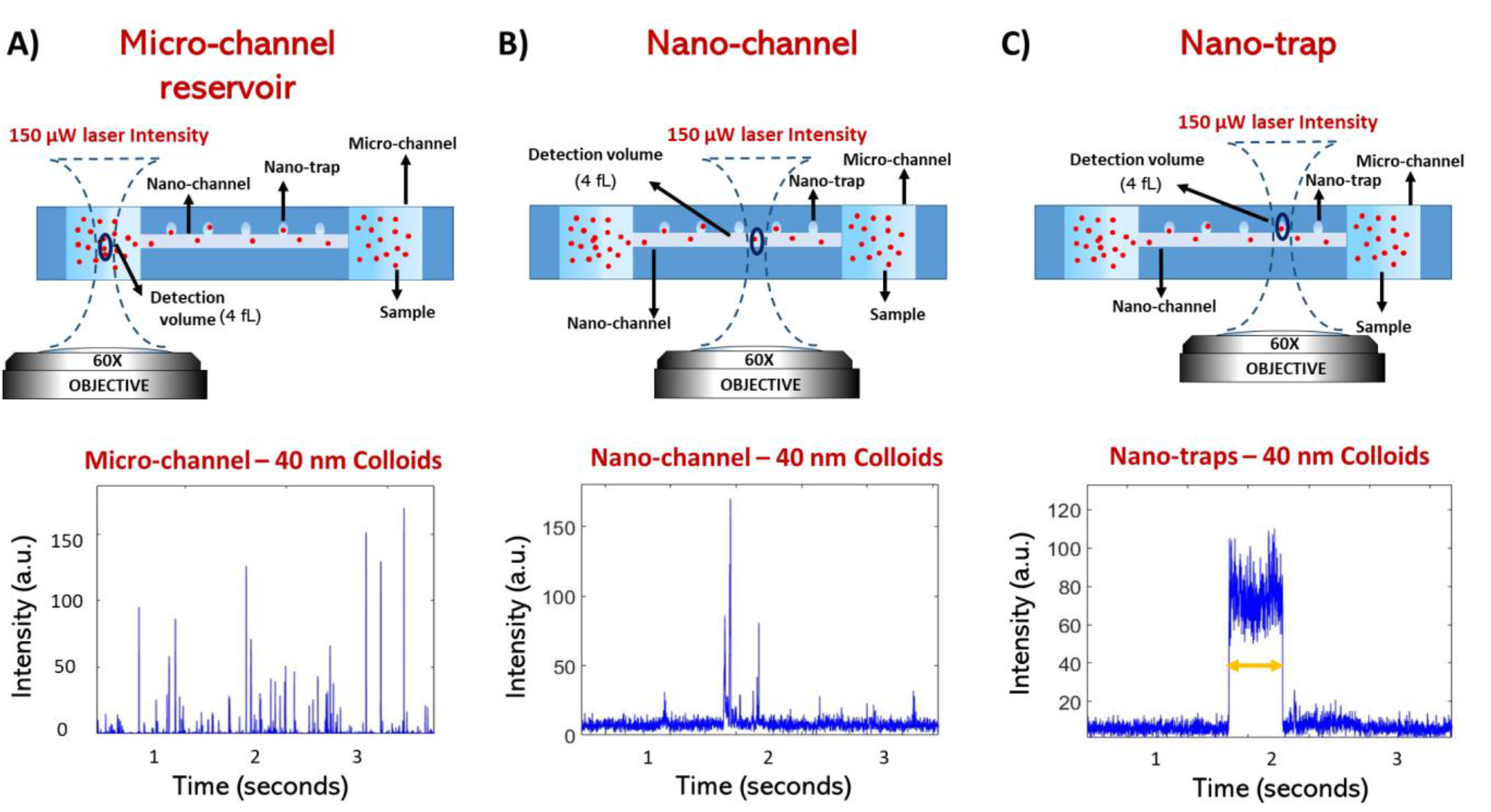
Single-molecule fluorescence detection in microfluidic reservoirs, nano-channels regions and under nano-trap confinement. **(A)** The confocal detection volume was placed into the microfluidic part of the device at the mid height of the channel (i.e., 13 μm above the glass cover slip). The diffusion of multiple particles at the same time through the confocal spot results in multiple fluorescence bursts as shown in the fluorescence burst time trace. **(B)** The confocal fluorescence burst detection volume was placed in the nano-channel region. Fluorescence data recorded in the nano-channel shows more rare events of fluorescent bursts, which implies that the probability of multiples particles crossing the detection volume is lowered by the nano-channel confinement. **(C)** The detection volume was placed into the centre of a nano-trap geometry. The fluorescence time trace data shows significantly increased residence time of single particles up to ten to hundreds of milliseconds under nano-trap confinement.

We first performed measurements on 40 nm fluorescent particles and compared burst detection under nano-trap confinement to residence times in the microfluidic reservoirs of the device and the nano-channel bridges. To this end, the confocal detection volume was placed in the respective region of the device, as illustrated in **Figure 5 (A)–(C)**. Within the microfluidic part of the device (**Figure 5 (A)**), multiple fluorescence burst signals are overlapping during the measurement and show various intensity levels, due to multiple particles being able to cross through the detection volume at the same time. The time regime of transition events is in the millisecond range. In a second measurement, the laser spot was placed inside a nano-channel, as shown in **Figure 5 (B)**, and confocal time traces were recorded. The number of fluorescence bursts was drastically reduced due to the single-molecule exclusion capabilities of the nano-channel, and just slightly increased detection times in comparison to measurements in the microfluidic channel were observed. Finally, we placed the confocal spot at the centre of a nano-trap. Nanoparticles in a single nano-trap geometry were recorded as shown in **Figure 5 (C)**. The time trace shown exemplifies the prolonged nature of fluorescence burst signals obtained within a nano-trap and is common amongst all species under study (**Supplementary Figure 3**).

Using the same nanofluidic geometry, we compared the behaviour of differently sized particles in the nano-traps. We performed experiments, as described before, with a series of nano-colloids and biomacromolecules, including 100 nm colloids, 40 nm colloids, α-synuclein oligomers (~9–14 nm), and 45 bp DNA (~15.3 nm length, estimated with 0.34 nm per bp, rod-like) [40][41] in deionized water. Our results show that the nano-traps increase the residence time of particles within the detection volume due to the additional local steric confinement. **Figure 6** shows a comparison of their mean residence times inside the nano-traps in relation to microfluidic channels. The time spent by the particle inside the laser spot depends on its diffusional properties and therefore on its size. In general, according to the Stokes-Einstein relation, the diffusion coefficient is defined as *D* = (*k_B_T*)/(6πη*R_H_*), where *R_H_* is the hydrodynamic radius, *k_B_* the Boltzmann constant, *T* the temperature and η the viscosity. This trend can be observed for confined and non-confined particles. Strikingly, comparing the nano-trap residence time to the microfluidic channel indicates an up to 5-fold increase of observation time within the confocal detection volume. This is expected because the walls limit the possibility of the molecule escaping from the laser’s field of view, as mentioned above. The Debye length can be assumed to be less than 100 nm [39] and should not be the major factor in the confinement presented here, but definitely needs to be considered when using smaller nanofluidic design dimensions instead. Enhancement of the residence time, once the particle is in the nano-trap, thus enables longer signal capture of a single particle. This opens-up the possibility for single-molecule metrology of biomolecules and colloids in solution over extended periods of time.

**Figure 6.**
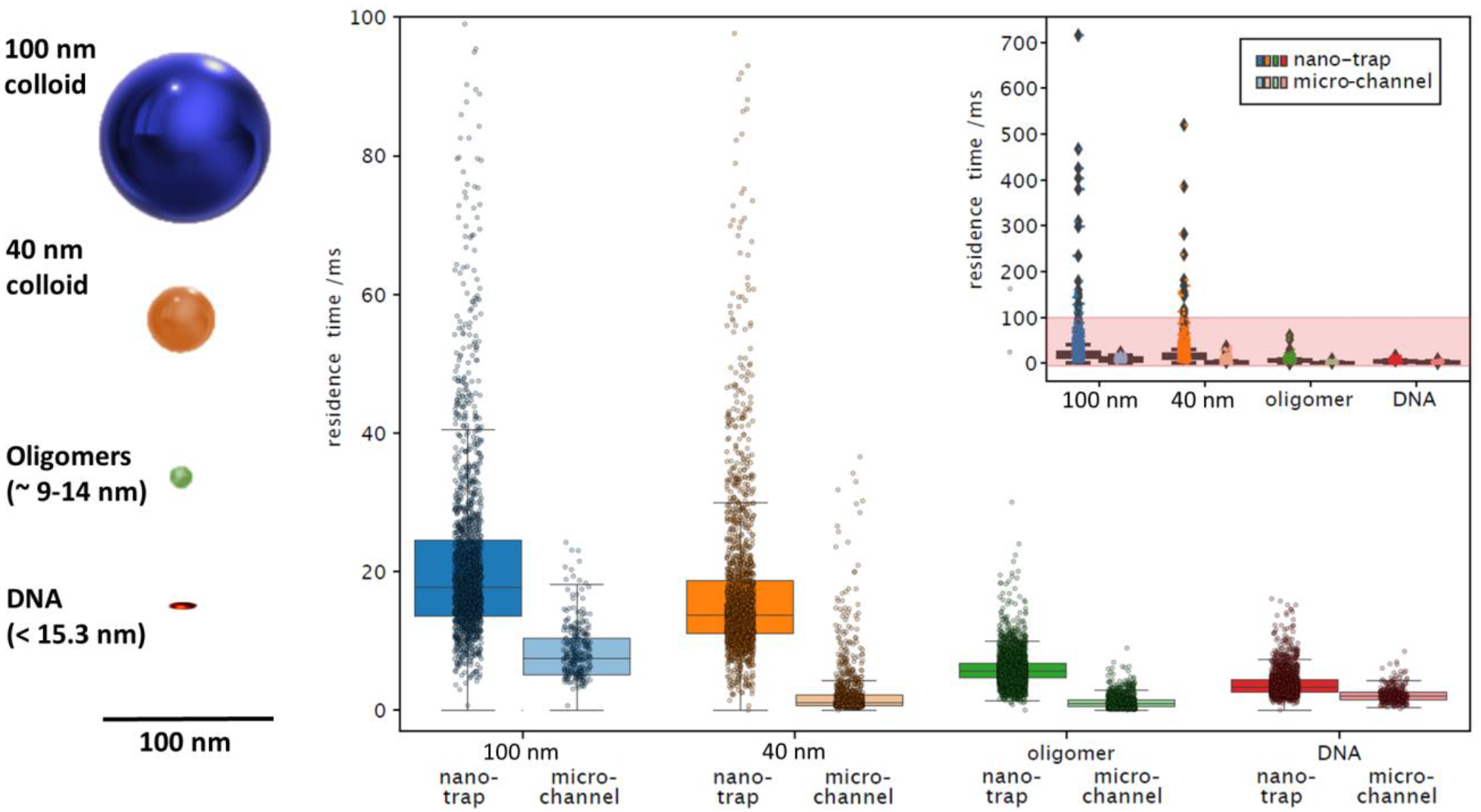
Residence time of specimen under nano-trap confinement. **(A)** Schematic illustration of the relative size difference of specimen probed. **(B)** Comparison between residence times for 100 nm colloids, 40 nm colloids, α-synuclein oligomers, and 45 bp DNA in micro-channel reservoirs and nano-trapping geometries. Residence time in nano-traps relative to the detection time in micro-channel reservoirs is increased by a factor of approximately 3- to 5-fold. The insert shows the existence of rare trapping events in the hundreds of millisecond range for colloidal particles, and up to tens of millisecond for oligomers and DNA.

## Conclusions

In this paper, we have demonstrated the use of hybrid 2PL for the fabrication of nano-traps written adjacent to nanofluidic and microfluidic channels and their usage for the study of colloidal nanoparticles and biomacromolecules at the single-molecule level. We have established conditions for the successful generation of a silicon master wafer with nanoconfinement geometries in a negative SU-8 photoresist by combining 2-photon direct laser writing with UV lithography. We imprinted nanofluidic devices from the silicon master wafer into PDMS to make functional nano-channels with adjacent nano-traps of 350 nm radius and 650 nm height, but also much smaller geometries, and structures below 100 nm in height, are possible (**Supplementary Figure 1**). Given the ease of fabrication, our approach can be readily adopted by laboratories with access to commercial or custom-built 2PL systems and allows for the fabrication and prototyping in a high-throughput and scalable manner as opposed to EBL and sequential clean room nanofabrication techniques.

To demonstrate the applicability of the nano-trapping devices developed herein for prolonged observation of single molecules, we used single-particle fluorescence burst detection to measure the residence time of polymer nanoparticles such as 100 nm and 40 nm colloids, and various biological relevant samples like α-synuclein oligomers and fluorescently labelled 45 bp DNA in nanofluidic confinement. Although our nano-trap geometry is orders of magnitude larger in comparison to the biological specimen under study, we observed a significant increase in residence times of the samples. All species analysed in the same trapping geometry showed up to 3- or 5-fold increase of observation time in a diffraction limited confocal detection volume. This finding is significant, as it opens-up the possibility to study and analyse biomacromolecules or biomolecular assemblies in solution without permanent surface immobilization for extended periods of time. It also allows longer observation of the same molecule for optical techniques that greatly benefit from higher photon counts such as FRET measurements at the single-molecule level.

Readout is not limited to single-particle fluorescence burst detection. As proof of concept, we also explored other fluorescence microscopy techniques (confocal imaging and TIRF microscopy) as alternative readout techniques to be used in combination with nanofluidics (**Supplementary Figure 4 and 5**). This gives laboratories guidance on how to use nano-trapping devices with their already available fluorescence microscopy equipment according to their needs and research applications. This highlights the versatility of the applications that can be envisaged with our nanofluidic device in conjunction with different optical modalities. We anticipate that the cost-effective and easy approach for fabrication of nanofluidic devices has the potential to find broad applicability in various applications in the nanobiotechnologies, biophysics, and clinical diagnostics.

Similar nanofluidic devices were previously established by Krishnan et al. for the geometry-induced electrostatic trapping of nano-colloids[18], where iSCAT provided a label-free readout method of gold nanoparticle and liposome residence times in the nanoconfinement. The silica-based devices were fabricated using RIE etching and involved several clean room fabrication steps - therefore are not easily prototyped by biological laboratories with limited access to nanofabrication facilities. An important step to make this technology more available to the research community was achieved by Gerspach et al. [42] who moulded electrostatic trapping devices in PDMS and measured the residence time of highly charged gold nanoparticles of 60 nm, 80 nm and 100 nm diameter in nano-pockets. Their experiments showed that confinement is highly dependent of the size ratio between the particle and the trap, which underlines the importance of flexible fabrication schemes that can adapt to the application accordingly.

By contrast, the method demonstrated in the present paper shows the advantage of a stationary chip design without external machinery to study a variety of biological specimen from colloids to oligomers and DNA molecules in confined space, without permanently immobilizing or perturbing these. EBL and RIE as the golden standards for the fabrication of silica trapping devices have higher lateral resolution than 2PL, but 2PL allows a more versatile integration of complex nanofluidic and nano-trapping geometries into microfluidic device platforms in the sub-micron regime.

Taken together, in this paper, we give a cost-effective and facile approach for the fabrication of nanofluidic devices to study single molecules in solution without permanent surface immobilization using hybrid 2-photon lithography. With our approach we envisage to facilitate nanoparticle trapping technology in biological and biomedical laboratories, paving the way for the use of photon-intensive spectroscopic techniques for applications related to protein misfolding disease, cancer research, and bionanotechnology.

## Methods

### Wafer preparation and development

SU-8 photoresist (Type 3025, Micro Resist Technology) was spin coated (Laurell technologies, WS-650) at 3000 rpm onto a 3-inch silicon wafer (MicroChemicals, Prime CZ-Si, thickness 381 +/- 20 μm, polished, p-type) to a height of 25 μm. The SU-8 coated wafer was soft baked and treated according to the protocol of the supplier of the photoresist. Microfluidic patterns from a custom-designed film mask (Microlithography) were then projected onto the wafer and the photoresist was exposed for 30 seconds with the UV-LED setup as described in Challa et al. [43]. The wafer was post baked at 95 °C so that the interfaces between exposed and unexposed regions become visible due to their change in refractive index, which assisted in alignment of the microstructures with the 2PL setup. After the nanostructures were written with 2PL and the wafer baked at 95 °C for 8 minutes. The wafer was developed using Propylene-glycol-monomethyl-ether-acetate (PGMEA) (Sigma-Aldrich) and subsequently given a second exposure with UV light for 30 seconds to make structures mechanically stable on the wafer before final rinsing of the structures with PGMEA and Isopropanol (IPA) (Sigma-Aldrich) [35]. A post-bake of 30 minutes at 95 °C on a hot plate was done at the end of the development process to increase mechanical stability of the nanostructures.

### 2-photon lithography

A custom-built 2PL setup was used to write the calibration patterns as well as the final nanofluidic master mould. A detailed description of the upright 2-photon lithography setup and its fabrication capabilities can be found in Vanderpoorten et al. [28]. Briefly, the system uses a femto-second fibre laser (Menlo System C Fiber 780 HP) modulated as the first diffraction order of an acousto-optic modulator (AA Optoelectronics). The beam is widened through a beam expander (Thorlabs, BE02-05-B) and led over a 90:10 R:T beamsplitter (BS028, Thorlabs) into a microscope objective vertically mounted above the sample. Reflected light is collected with a tube lens (Thorlabs AC 254-100-A-ML, BBAR coating A OM 31 400-700 nm, f = 100.0 mm) onto a camera (μEye ML, Industry camera, USB 3.0). An optical electro-mechanical shutter (Thorlabs, SHB1) is mounted in front of the camera to protect it during high power laser writing. Through an additional 30:70 (R:T) beam splitter (BS019, Thorlabs) in the camera detection arm, a white LED (Thorlabs, MCWHL5) allows non-polymerizing inspection of the sample in wide field. A 3-inch wafer coated with pre-baked SU-8 (25 μm thickness) was immobilised on a PI Nanocube (P-611.3S, Physikalische Instrumente) mounted on two perpendicular stacked motorised linear-precision stages (M-404.2PD, Physikalische Instrumente, Ball screw, 80 mm wide, ActiveDrive). Immersion Oil (Cargille laboratories, LDF, Code 387) was added onto the SU-8 layer before bringing the oil immersion objective (Leica, 63x, PL APO, 1.40 NA) manually in close proximity to the wafer surface. The oil used here showed no reaction with unpolymerised SU-8 photo resin and facilitates easy and scalable two-photon printing. Custom-written software then automatically focusses on the wafer surface, corrects for tilt and coordinates the interplay of piezo, translational stages and laser power modulation to write the intended patterns. The laser beam intensity of the writing beam was directly measured after the acousto-optic modulator using a power meter (Thorlabs, S310C, thermal power head). To prevent exposure of the resin during the focussing process, the laser power was kept below the polymerization threshold, but high enough to be detected on the system’s camera. The full travel range of the Nanocube of 100 μm x 100 μm was used to write a calibration array of lines and dots. Then the motorized stages were used to displace the piezo scanning areas and write a new pattern (e.g., 300 μm displacement, positional precision = 1 μm) with adapted parameters. The positioning repeatability of the piezo actor (Nanocube) was below 10 nm according of the manufacturer and is key for automated focussing and reliable nanofabrication. For 2-photon-writing in the microfluidic master, we used a white light LED to first place the laser focus in between the two micro-channels and then started the automated laser writing process. The system uses the autofocus function each time it adds another nanofluidic array. This allows step wise but precise addition of nanofluidic features on the wafer scale.

### Correlative scanning electron microscopy and atomic force microscopy imaging

After the development process of the 2-photon written calibration assay, the wafer was manually cut into smaller dimensions to allow easier sample handling. Imprints of the master wafer were taken following conventional soft lithography protocols PDMS (Sylgard 184) with 10:1 curing agent ratio. After PDMS curation, the area of interest was cut out using a surgical scalpel. The PDMS imprint was coated with 10 nm platinum (Quorum Technologies Q150T ES Turbo-Pumped Sputter Coater/Carbon Coater) and imaged using a commercial SEM (TESCAN MIRA3 FEG-SEM). The original SU-8 features were coated with a layer of 10 nm platinum as well and imaged on the same SEM in order to compare the imprinted features with the original moulds. The final nanofluidic PDMS device imprint was imaged following the same procedures and imaged on the same microscope. AFM was conducted on the calibration sample using a Park Systems NX10 AFM. According to previous findings by Cabrera et al. [44] the PDMS surface roughness can be assumed to be below 5 nm, which should therefore not influence the steric trapping behaviour significantly.

### Profilometer measurements of nano-traps

The 2-photon written nanofluidic master wafer was cleaned using pressurised air and placed in a profilometer (KLA Corporation, Tencor P-6) for height measurements of nano-channels and nano-traps. Using the integrated microscope of the system, the scan direction was aligned along the centre of a nano-trapping array located between the two microfluidic reservoirs. The sample was scanned at a speed of 2.00 μm/s, with a height scan rate of 500 Hz and a force of 0.5 mg applied using a 2.00 μm (diameter) tip.

### Single-molecule confocal measurements

Single-molecule fluorescence measurements were performed on a custom-built single-molecule confocal microscope. Nanofluidic PDMS–silica devices were secured to a motorised microscope stage (Applied Scientific Instrumentation, PZ-2000FT). The sample was excited using a 488 nm wavelength laser (Cobolt 06-MLD, 200 mW diode laser, Cobolt), which was directed to the back aperture of a 60X-magnification water-immersion objective (CFI Plan Apochromat WI 60x, NA 1.2, Nikon) using a single-mode optical fibre (P3-488PM-FC-1, Thorlabs) and an achromatic fibre collimator (60FC-L-4-M100S-26, Schäfter/Kirchhoff GmbH). The laser intensity at the back aperture of the objective was adjusted to 150 μW. The laser beam exiting the optical fibre was reflected by a dichroic mirror (Di03-R488/561, Semrock), directed to the objective and focussed into the chip to a diffraction-limited confocal spot. The motorised stage was used to position the confocal spot within the chip. The emitted light from the sample was collected through the same objective and dichroic mirror and then passed through a 30 μm pinhole (Thorlabs) to remove any out-of-focus light. The emitted photons were filtered through a band-pass filter (FF01-520/35-25, Semrock) and then focussed onto an avalanche photodiode (APD, SPCM-14, PerkinElmer Optoelectronics) connected to a TimeHarp260 time-correlated single-photon counting unit (PicoQuant). Photon time traces were recorded using the SymPhoTime 64 software package (Picoquant) with a binning time of 1 ms.

### Preparation of labelled α-synuclein oligomers

The N122C variant of α-synuclein was purified into phosphate buffered saline (PBS) pH 7.4 as described previously [45], with the addition of 3 mM DTT to all buffers to prevent dimerization. Following removal of DTT from the purified monomers by a PD10 desalting column packed with Sephadex G25 matrix (GE Healthcare), the protein was incubated with a 1.5-fold molar excess of Alexa488 with a maleimide linker (ThermoFisher Scientific) (overnight, 4 °C on a rolling system). In order to remove the free dye, the mixture was subsequently subjected to size exclusion chromatography using a Superdex 200 16/600 (GE Healthcare) and eluted in PBS pH 7.4 at 20 °C. Protein fractions were pooled, and Alexa488 labelled α-synuclein concentration estimated by dye absorbance, assuming 1:1 dye:protein stoichiometry (72 000 L/mol cm at 495 nm). Stable α-synuclein oligomers were formed from Alexa488 labelled monomers, as previously described [46]. Briefly, monomeric α-synuclein was lyophilised in Milli-Q water and resuspended in PBS pH 7.4 at a concentration of 12 mg/m. Following incubation (37 °C, 20-24 h), the samples were ultracentrifuged (1h, 288’000 x g) (Optima TLX Ultracentrifuge, Beckman Coulter, TLA-120.2 Beckman rotor) to remove large aggregates. Monomeric protein was removed by multiple filtration steps through 100 kDa concentrating filters. The oligomer concentration was estimated based on the dye absorbance (72’000 L/mol cm at 495 nm).

### Sample and device preparation for single molecule experiments

100 nm and 40 nm fluorescent colloids (FluoSpheres) were purchased from ThermoFisher. α-synuclein oligomers were prepared as described above. Double-stranded DNA was prepared from two single-stranded DNA oligonucleotides by thermal annealing. Oligonucleotides were synthesized and labelled by Biomers. The sequences were: 5’-GCC TTA **T**TT TCA CTC TTT CCT TTC TTC TTC TCT CTT TTT TTC CCG-3’ (top strand) and 5’-CGG GAA AAA AAG AGA GAA GAA GAA AGG AAA GAG TGA AAA TAA GGC-3’ (bottom strand); the top strand was labelled with Atto488 at the thymidine at position 7, shown in bold type.

Micro-/nanofluidic devices were moulded from the fabricated SU-8 master via soft lithography using PDMS (Sylgard 184; with 10:1 curing agent ratio). After baking, inlets were added using surgical punchers and plasma bonded to coverslip glasses (Menzel coverslips, Grade H1.5). The surface of the coverslips and the PDMS were plasma treated, and afterwards manually pressed on top of each other. Devices were used directly after the plasma bonding step to use their remaining surface hydrophilicity for easier filling of the devices. Before the experiments, the chips were filled by pipetting equal amounts of diluted sample solutions into the inlet areas and equilibrated for 20 minutes.

## Acknowledgements

The work was funded by the Horizon 2020 programme through 766972-FET-OPEN-NANOPHLOW (TPJK). The research leading to these results has further received funding from the European Research Council under the European Union’s Horizon 2020 Framework Programme through the Marie Sklodowska-Curie grant MicroSPARK (agreement n° 841466; GK), the Herchel Smith Funds (GK), and the Wolfson College Junior Research Fellowship (G.K.). This work was also supported by the Engineering and Physical Sciences Research Council [grant numbers EP/L015889/1]. The authors would also like to thank the NanoDTC for additional funding and the Maxwell Community for scientific support.

## Author contributions statement

OV and ANB fabricated nanofluidic masters and chips using two-photon lithography. ANB characterised the calibration assays using SEM and AFM. OV conducted profilometer measurements on nanofluidic master wafer. GK built the confocal burst detection setup and conducted the fluorescence burst experiments of colloids, oligomers, and DNA in nano-trap devices and micro-channels. OV and ANB imaged trapping events of colloidal particles using confocal microscopy. PKC, ZT, ANB and OV conducted TIRF microscopy measurements of colloidal particles. PKC and ZT also conducted trapping experiments using conventional UV-lithography in an early stage of the project, which helped form the content of this paper. RJ contributed with data analysis and wrote software for the residence time measurements of the fluorescence burst data. QP improved the control software of the two-photon system which allowed initial test runs and to conduct the fabrication assay. CX prepared and purified oligomer samples used for all the experiments. OV, GK, and ANB wrote the paper. All authors provided input into the manuscript.

