## Supplementary Information for "2-photon-fabricated nano-fluidic traps for extended detection of single macromolecules and colloids in solution"

### Supplementary Figure 1

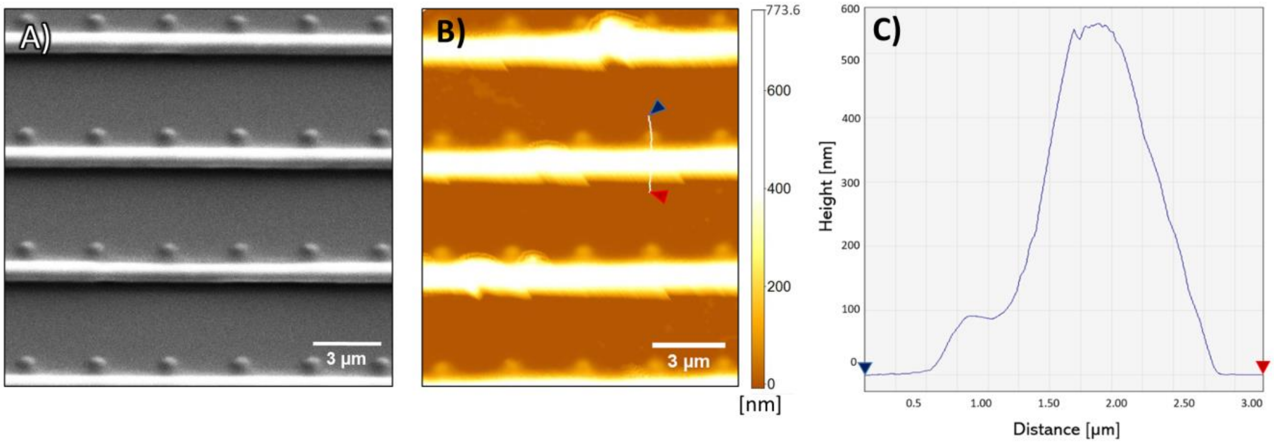


**Supplementary Figure 1. Correlative SEM and AFM analysis of nano-channels and nano-traps. (A)** SEM micrograph of nano-channel and nano-trap structures written with two photon lithography in SU-8 photoresist by ascending the voxel into the wafer. **(B)** AFM imaging reveals a height of 100 nm for nano-traps and 550 nm for nano-channels. **(C)** AFM line scan profile of nano-channel and nano-trap structures (as indicated in (B)).

**Supplementary Figure 2**

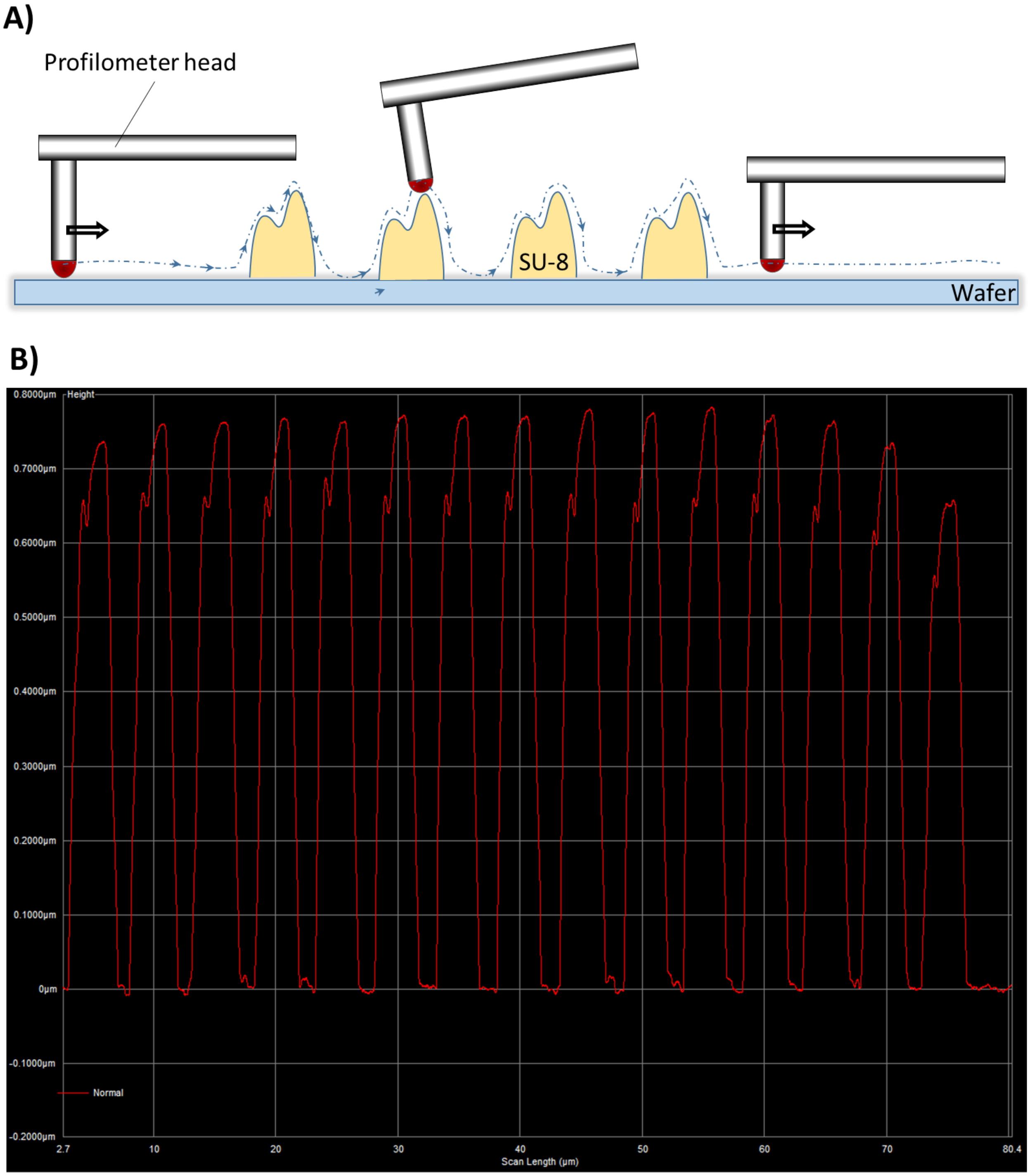


**Supplementary Figure 2. Profilometer measurement of nano-channel array with integrated nano-traps.** The measurement was done on the device used for single-molecule experiments. **(A)** Shows schematically how the profilometer was used to scan the sample. **(B)** The line scan shows the height profile of several nano-traps with heights of approx. 650 nm connected to nano-channels with heights of 750 nm. Scan speed = 2.00 µm/s, force = 0.5 mg, 2.00 µm (diameter) tip.

### Supplementary Figure 3

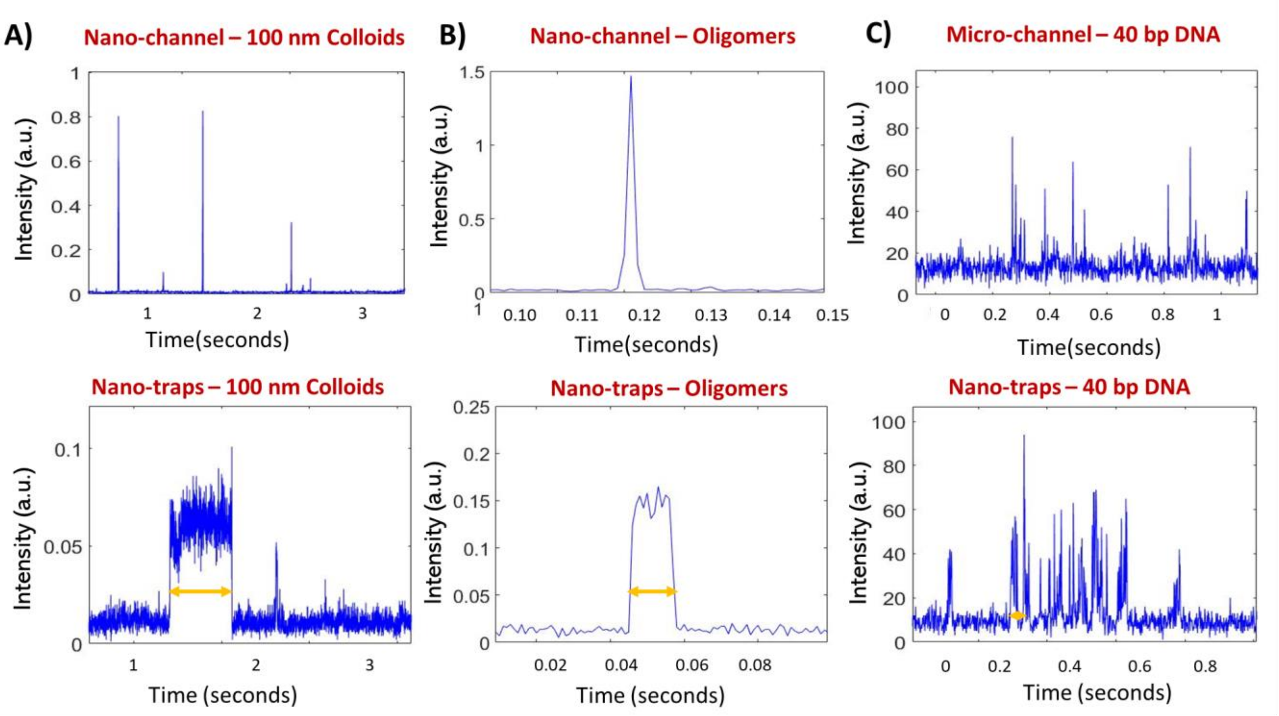


**Supplementary Figure 3. Single-molecule fluorescence detection of 100 nm colloids, α-synuclein oligomers and 40 bp DNA in nano-channels regions and under nano-trap confinement. (A)** Measurement of fluorescence burst traces in nano-channel and nano-trap confinement for 100 nm colloids. **(B)** Measurement of fluorescence burst traces in nano-channel and nano-trap confinement for αS oligomers. **(C)** Measurement of fluorescence burst traces in micro-channel and nano-trap confinement for 45 bp DNA.

**Supplementary Figure 4**

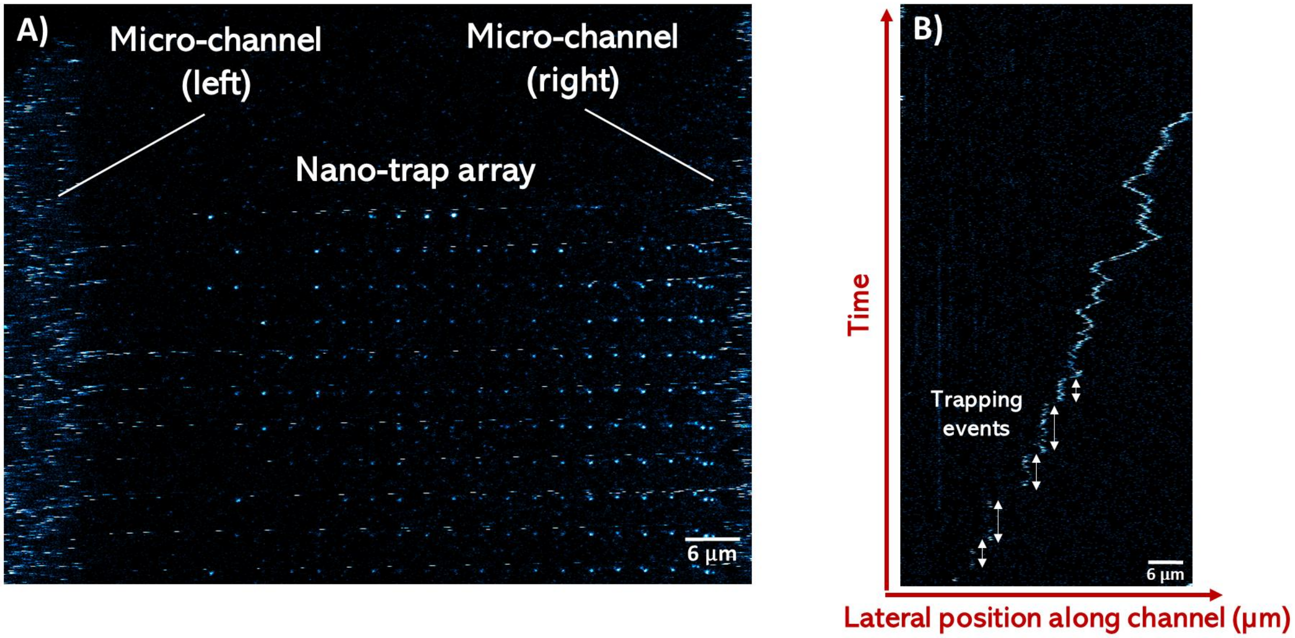


**Supplementary Figure 4. Observation of colloidal particle trapping in nanofluidic trapping device by confocal line scanning fluorescence microscopy. (A)** The nano-fluidic device was filled with 100 nm colloids and images using confocal line scanning microscopy. Colloids appear as fluorescent streaks within the microfluidic regions and nano-channel of the device due to the rapid movement of the sample. Conversely, particles in nano-traps are visible as round objects, demonstrating that colloid are spatially confined in the nano-trap cavities. **(B)** Kymograph analysis of particle movement within nano-channels and nano-trap cavities. Confinement in the nano-trap geometry is indicated by white arrows and evident as vertical lines in the space-time plots. Line scan time was 12.5 ms (frame rate of 81 Hz) at a pixel dwell time of 25 µs at 200 nm pixel.

**Observation of colloidal particle trapping in nanofluidic trapping devices using confocal fluorescence microscopy.** The combined nano-fluidic device was mounted onto an Abberior RESOLFT confocal imaging setup and fluorescent 100 nm-sized colloids at nanomolar concentration were flushed from the inlets into both micro-channels by manual pipetting. Imaging was performed with a 489 nm excitation laser beam and a 520/10 nm emission bandpass filter for detection. A 100x Olympus objective was used for imaging, which resulted in a field of view of 80 µm x 80 µm, which is large enough to image parts of the micro-channels, the nano-channel entries and the nano-traps within one scanning frame as indicated in **Supplementary Figure 4 (A)**. Particles that move during the acquisition result in stripes in a confocal line scanning image, whereas confined particles remain in place and appear as point-like objects. In **Supplementary Figure 4 (A)**, the microfluidic reservoirs can be seen on the left and right of the image, where streaks are indicative of particle movement of the colloidal suspension. Similarly, within nano-channels, particles appear as streaks due to their movement along the nano-channels. Conversely, within the nano-trap arrays, particles appear as spherical, point-like objects. Kymograph analysis (**Supplementary Figure 4 (B)**), performed by repeatedly scanning over a single nano-channel, indicate that particles enter and leave the nano-traps after being confined in the traps for a certain period of time (vertical lines, represented with arrows).

### Supplementary Figure 5

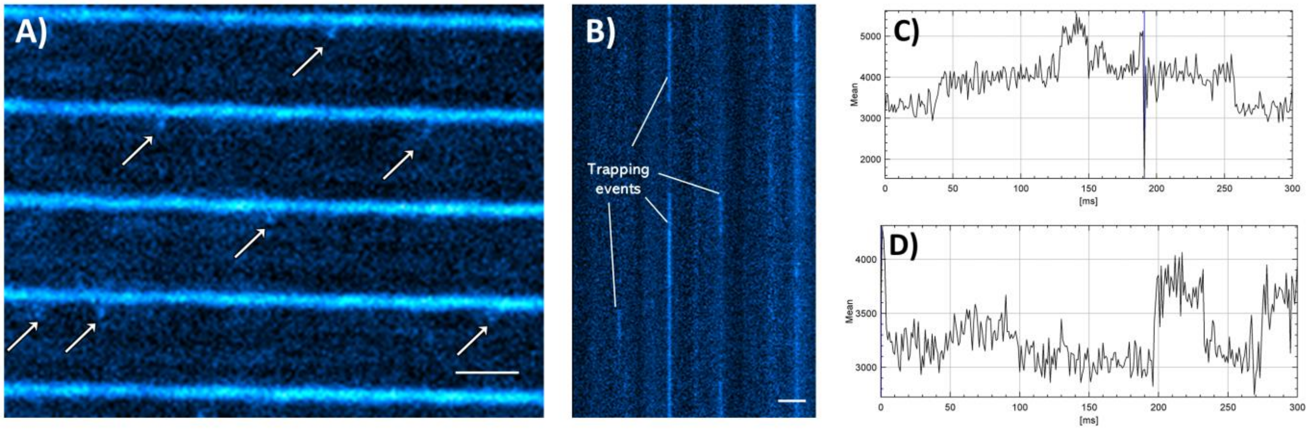


**Supplementary Figure 5. Observation of nanoparticle confinement within nano-traps using TIRF microscopy. (A)** Fluorescence image of 40 nm colloids within nanofluidic device. Arrows indicate the presence of particles in the nano-traps adjacent to nano-channels filled with colloidal suspension. Frame rate: 1 kHz. **(B)** Kymograph showing trapping events of 40 nm colloids in nano-traps. The length of the plot in the time domain is 300 ms. Data was processed using FIJI. **(C, D)** Intensity plot along a single nano-trap shows discrete intensity steps due to single-particle trapping events. Scale bars = 3 µm.

**Observation of colloidal nanoparticle confinement in nano-trapping arrays using TIRF microscopy.** Nano-trap arrays with dimensions of 75 µm x 75 µm were imaged using TIRF microscopy with a kHz framerate camera. In this experiment, the devices were filled with 40 nm fluorescent colloids. Images were acquired with an Evolve Delta EMCCD camera, using a fibre-coupled 485 nm laser (Picoquant) as the excitation source and a GFP filter set on a commercial Nikon Ti-E inverted microscope installed with a motorised TIRF module at the back of the microscope. **Supplementary Figure 5 (A)** shows an exemplary frame of the acquired data from imaging 40 nm colloids in a nanofluidic device. The nano-channels exhibit a continuous fluorescent signal due to the fast colloidal movements and the relatively high particle density that blurs the image and renders single particle observation impossible. However, emitters appearing and disappearing below the nano-channels within nano-trap cavities indicate the transient confinement of single particles. Similarly, as in the confocal scanning microscopy experiment above, kymographs were generated from the image stacks to evaluate trapping events in the time domain. For illustration purposes, a kymograph is plotted in **Supplementary Figure 5 (B)**, with single-particle trapping events being visible as vertical lines. In **Supplementary Figure 5 (C, D)**, we found quantized intensity bursts at discrete fluorescence intensity levels, indicating single-particle trapping events.
